# Large-scale tandem mass spectrum clustering using fast nearest neighbor searching

**DOI:** 10.1101/2021.02.05.429957

**Authors:** Wout Bittremieux, Kris Laukens, William Stafford Noble, Pieter C. Dorrestein

## Abstract

**Rationale:** Advanced algorithmic solutions are necessary to process the ever increasing amounts of mass spectrometry data that is being generated. Here we describe the *falcon* spectrum clustering tool for efficient clustering of millions of MS/MS spectra.

**Methods:** *falcon* succeeds in efficiently clustering large amounts of mass spectral data using advanced techniques for fast spectrum similarity searching. First, high-resolution spectra are binned and converted to low-dimensional vectors using feature hashing. Next, the spectrum vectors are used to construct nearest neighbor indexes for fast similarity searching. The nearest neighbor indexes are used to efficiently compute a sparse pairwise distance matrix without having to exhaustively perform all pairwise spectrum comparisons within the relevant precursor mass tolerance. Finally, density-based clustering is performed to group similar spectra into clusters.

**Results:** Several state-of-the-art spectrum clustering tools were evaluated using a large draft human proteome dataset consisting of 25 million spectra, indicating that alternative tools produce clustering results with different characteristics. Notably, *falcon* generates larger highly pure clusters than alternative tools, leading to a larger reduction in data volume without the loss of relevant information for more efficient downstream processing.

**Conclusions:** *falcon* is a highly efficient spectrum clustering tool. It is publicly available as open source under the permissive BSD license at https://github.com/bittremieux/falcon.

## 1 Introduction

To obtain a comprehensive view of an organism’s proteome, modern shotgun proteomics experiments generate thousands [21, 46] to tens of thousands [18, 19] of tandem mass spectrometry (MS/MS) runs, with tens of thousands of MS/MS spectra acquired during each individual run. Besides the significant efforts to acquire such large amounts of spectral data, processing these large data volumes poses a computational challenge that requires efficient algorithmic solutions.

Typically, the MS/MS spectra are processed using a sequence database search engine to derive their peptide and protein identities [35]. Alternatively, rather than having to search all of the raw spectra, as a preprocessing step spectrum clustering can be used to reduce the data volume [9, 12, 13, 38, 40]. Spectrum clustering groups highly similar spectra, after which each cluster can be represented by a single consensus spectrum. In this fashion a data reduction can be achieved because only the cluster representatives need to be processed. Additionally, because consensus spectra can have a higher signal-to-noise ratio than the raw spectra and because low-quality, unclustered spectra can be filtered out, the clustering approach can boost the sensitivity of the subsequent identification procedure. Furthermore, repository-scale clustering can be used to automatically generate comprehensive and high-quality spectral libraries in a data-driven fashion without having to rely on synthetic samples [12, 42], and the clustering results can be analyzed to gain deeper insights into the nature of repeatedly observed yet unidentified spectra [13].

Several spectrum clustering tools have been introduced, including MS-Cluster [9], spectra-cluster [12, 13], MaRaCluster [38], and msCRUSH [40]. In general, a clustering algorithm consists of several components: (i) a similarity measure to perform pairwise spectrum comparisons, (ii) a clustering method to group similar spectra, and (iii) optional optimizations to improve its computational efficiency. MS-Cluster [9] uses the cosine similarity as similarity measure. It obtains an approximate hierarchical clustering result by merging spectra that exceed an iteratively decreasing similarity threshold in a greedy fashion (rather than always merging the most similar spectra, as in standard hierarchical clustering). To avoid unnecessary similarity calculations only pairs of spectra that share at least one peak among their five most intense peaks are compared to each other. The spectra-cluster approach was originally developed as a reimplementation of MS-Cluster, with some refinements to improve the cluster quality [12]. A highly parallel implementation was subsequently developed [13] to efficiently cluster large amounts of public data available in the PRoteomics IDEntifications (PRIDE) database [31]. Similar to MS-Cluster, spectra-cluster uses an iterative greedy approach to merge similar spectra. In contrast, instead of the cosine similarity a probabilistic scoring scheme [7] was adopted as similarity measure [13]. MaRaCluster [38] uses a specialized similarity measure that relies on the rarity of fragment peaks to compare MS/MS spectra. Based on the intuition that peaks shared by only a few spectra offer more evidence than peaks shared by a large number of spectra, relative to a background frequency of fragment peaks with specific *m/z* values, matches of highly frequent fragment peaks contribute less to the spectrum similarity than matches of rare peaks. Next, MaRaCluster uses hierarchical clustering with complete linkage to group similar spectra in clusters. Finally, msCRUSH [40] is a fast spectrum clustering tool based on locality-sensitive hashing [11]. By efficiently hashing similar spectra to identical buckets, unnecessary pairwise spectrum comparisons can be avoided. Next, within each bucket a similar greedy spectrum merging strategy is performed as employed by MS-Cluster and spectra-cluster.

Because each of these spectrum clustering tools use different spectrum similarity measures, clustering methods, and computational optimizations, their clustering results and computational performance will exhibit different characteristics. Here we introduce *falcon*, a fast spectrum clustering approach. By making use of advanced algorithmic techniques, *falcon* is optimized for highly efficient spectrum clustering. It uses feature hashing to convert high-resolution MS/MS spectra to low-dimensional vectors [4], in combination with efficient nearest neighbor searching in the vector metric space using the cosine similarity [5]. Next, spectra are grouped into clusters by density-based clustering [8]. We compare *falcon* to the state-of-the-art clustering tools MaRaCluster, MS-Cluster, msCRUSH, and spectra-cluster in terms of clustering quality and runtime, and show that it succeeds in efficiently clustering large amounts of spectral data. *falcon* is freely available as open source under the permissive BSD license at https://github.com/bittremieux/falcon.

## 2 Methods

### 2.1 Spectrum preprocessing

The spectra are preprocessed by removing peaks corresponding to the precursor ion and low-intensity noise peaks, and, if applicable, spectra are further restricted to their 50 most intense peaks. Low-quality spectra that have fewer than five peaks remaining or with a mass range between their minimum and maximum peak less than 250 Da after peak removal are discarded. Finally, peak intensities are square root transformed to de-emphasize overly dominant peaks [25].

### 2.2 Feature hashing to convert high-resolution spectra to low-dimensional vectors

To build a nearest neighbor index for efficient spectrum similarity searching, spectra need to be vectorized to represent them as points in a multidimensional space. MS/MS spectra typically contain dozens to hundreds of peaks, whose *m/z* values are measured at a resolution in the order of 1/100 *m/z*. As such, a straightforward approach to convert spectra to vectors by dividing the mass range into small bins and assigning each peak’s intensity to the corresponding bin would result in extremely high-dimensional, sparse vectors that are not suitable for efficient nearest neighbor searching due to the curse of dimensionality [2]. Alternatively, larger mass bins can be used to reduce the vectors’ dimensionality and sparsity. However, because such larger mass bins considerably exceed the fragment mass tolerance when dealing with high-resolution spectra, multiple distinct fragments can get merged into the same mass bin. This merging leads to an overestimation of the spectrum similarity when comparing two spectra to each other using their vector representations due to spurious matches between fragments.

Instead, a feature hashing scheme [45] is used to convert high-resolution MS/MS spectra to lowdimensional vectors while closely capturing their fine-grained mass resolution. The following two-step procedure is used to convert a high-resolution MS/MS spectrum to a vector (figure 1A) [4]:

1. Convert the spectrum to a sparse vector using small mass bins to tightly capture fragment masses.
2. Hash the sparse, high-dimensional, vector to a lower-dimensional vector by using a hash function to map the mass bins separately to a small number of hash bins.

**Figure 1:**
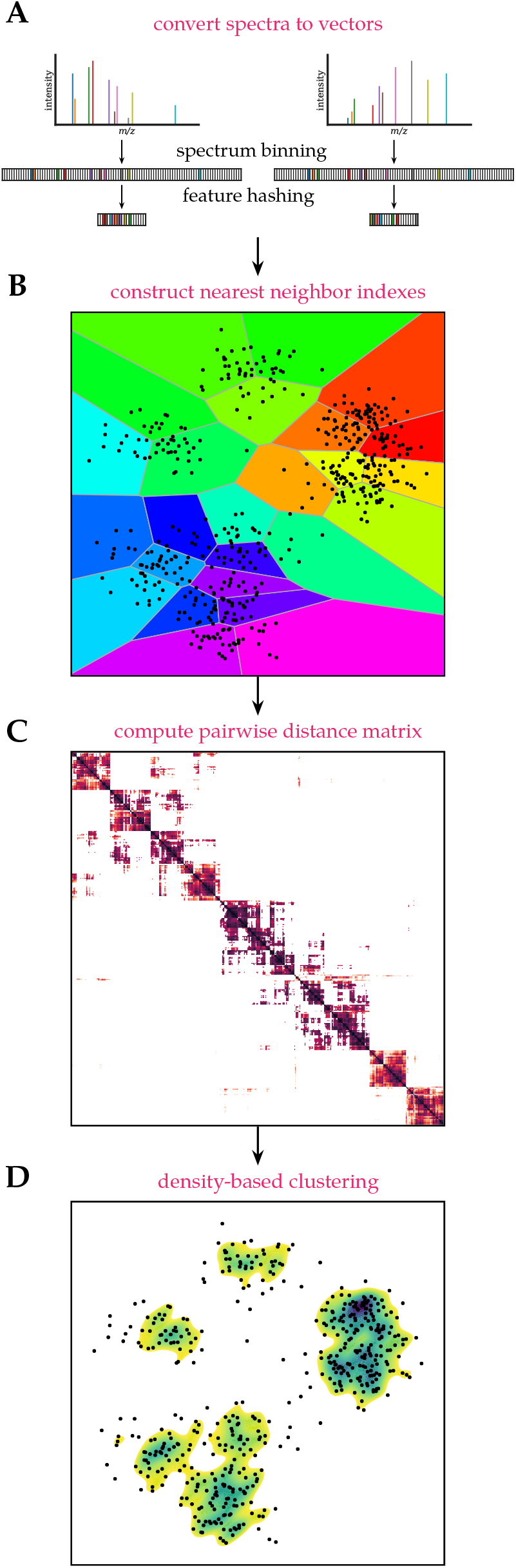
*falcon* spectrum clustering workflow. **(A)** High-resolution MS/MS spectra are converted to low-dimensional vectors using feature hashing. **(B)** Vectors are split into intervals based on the precursor *m/z* of the corresponding spectra to construct nearest neighbor indexes for highly efficient spectrum comparison. **(C)** A sparse pairwise distance matrix is computed by retrieving similarities to neighboring spectra using the nearest neighbor indexes. **(D)** Density-based clustering using the pairwise distance matrix is performed to find spectrum clusters.

More precisely, let *h*: 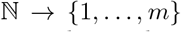 be a random hash function. Then h can be used to convert a vector *x* = (*x*_1_,…, *x_n_*) to a vector 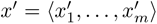, with *m* ≪ *n:*

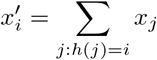

As hash function h the MurmurHash3 algorithm [1], a popular non-cryptographic hash function, is used.

It can be proven that under moderate assumptions feature hashing approximately conserves the Euclidean norm [10], and hence, the cosine similarity between hashed vectors can be used to approximate the similarity between the original, highdimensional vectors and spectra.

Note that this feature hashing procedure operates on each mass bin individually. In contrast, during locality-sensitive hashing, for example, as employed by msCRUSH [40], entire spectra are hashed as a single entity.

### 2.3 Efficient density-based clustering using nearest neighbor searching

Nearest neighbor searching is used to process large search spaces for efficient spectrum clustering [5]. Per precursor charge the MS/MS spectra are partitioned into 1 *m/z* buckets based on their precursor mass and converted to vectors as described previously. Next, the spectrum vectors in each bucket are partitioned into data subspaces to create a Voronoi diagram (figure 1B). The Voronoi diagram is encoded by an inverted index, with each Voronoi cell defined by a single vector, determined using k-means clustering to find a user-specified number of representative vectors, and all vectors are assigned to their nearest representative vector. This inverted index can then be used for efficient similarity searching. Instead of having to compare all spectrum vectors to all other vectors in the bucket to find their nearest neighbors, after mapping the vectors to their Voronoi cells they only need to be compared to the limited number of vectors therein.

The accuracy and speed of similarity searching is governed by two hyperparameters: the number of Voronoi cells to use during construction of the inverted index and the number of neighboring cells to explore during searching. Using a greater number of Voronoi cells achieves a more fine-grained partitioning of the data space, and exploring more cells during searching decreases the chance of missing a nearest neighbor in the high-dimensional space. In practice, for *m/z* buckets that contain fewer than 100 spectra a brute-force search is used. For larger *m/z* buckets the number of Voronoi cells is dynamically set based on the number of spectra in the bucket *N*. For buckets that consist of up to one million spectra 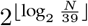 Voronoi cells are used, for buckets that consist of up to ten million spectra 2^16^ Voronoi cells are used, for buckets that consist of up to one hundred million spectra 2^18^ Voronoi cells are used, and for larger buckets 2^20^ Voronoi cells are used. During searching maximum 32 neighboring Voronoi cells per query vector are explored.

This efficient similarity searching is used to construct a sparse pairwise distance matrix that contains the cosine distances between each spectrum and a limited number of its nearest neighbors, additionally filtered using a precursor mass tolerance (figure 1C). Besides being able to retrieve the nearest neighbors highly efficiently, not having to compute and store all pairwise similarities also avoids extreme memory requirements.

Next, the pairwise distance matrix is used to cluster the data using the DBSCAN algorithm (figure 1D) [8, 33]. Briefly, if a given number of spectra are close to each other and form a dense data subspace, with closeness defined relative to a user-specified distance threshold, they will be grouped in clusters. An important advantage of DBSCAN is that the number of clusters is not required to be known in advance. Instead, it is able to find clusters in dense regions, whereas spectra in low-density regions, without a sufficient number of close neighbors, will be marked as noise. Additionally, DBSCAN is scalable: using a sparse pairwise distance matrix as input it can effortlessly process millions to billions of data points.

A disadvantage of this clustering approach, however, is that despite using a precursor mass filter during construction of the pairwise distance matrix, spectra within a single cluster can still exceed the precursor mass tolerance if they are connected by another spectrum with an intermediate precursor mass [33]. To avoid such false positives, the clusters reported by DBSCAN are postprocessed by hierarchical clustering with maximum linkage of the cluster members’ precursor masses. In this fashion, some clusters are split into smaller, coherent clusters so that none of the spectra in a single cluster have a pairwise precursor mass difference that exceeds the precursor mass tolerance.

### 2.4 Data

Spectrum clustering was performed on the human draft proteome dataset by Kim et al. [21]. This dataset aims to cover the whole human proteome and consists of 30 human samples in 2212 raw files, corresponding to 25 million MS/MS spectra. For full details on the sample preparation and acquisition see the original publication by Kim et al. [21]. Raw files were downloaded from PRIDE [31] (project PXD000561) and converted to MGF files using ThermoRawFileParser (version 1.2.3) [16].

Spectrum identifications were downloaded from MassIVE reanalysis RMSV000000091.3. These identifications were obtained via automatic reanalysis of public data on MassIVE using MS-GF+ [22]. Spectra were searched against the UniProtKB/Swiss-Prot human reference proteome (downloaded 2016/05/23) [6] augmented with common contaminants. Search settings included a 50 ppm precursor mass tolerance, trypsin cleavage with maximum one non-enzymatic peptide terminus, and cysteine carbamidomethylation as a static modification. Methionine oxidation, formation of pyroglutamate from N-terminal glutamine, N-terminal carbamylation, N-terminal acetylation, and deamidation of asparagine and glutamine were specified as variable modifications, with maximum one modification per peptide. Peptide-spectrum matches (PSMs) were filtered at 1 % false discovery rate, resulting in 10 487 235 spectrum identifications.

All clustering results are available on Zenodo at doi:10.5281/zenodo.4721496.

### 2.5 Clustering evaluation

For evaluation purposes, only MS/MS spectra with the common precursor charges 2 and 3 are considered. Valid clusters are required to consist of minimum two spectra. *falcon* explicitly designates non-clustered spectra as noise points with cluster label “-1”. In contrast, MaRaCluster, MS-Cluster, msCRUSH, and spectra-cluster report singleton clusters consisting of single spectra with unique cluster labels. To evaluate the cluster quality in a consistent fashion, such clusters are postprocessed and labeled as noise as well.

The following evaluation measures are used to assess cluster quality:

**Clustered spectra** The number of spectra in non-noise clusters divided by the total number of spectra.
**Incorrectly clustered spectra** The number of incorrectly clustered spectra in non-noise clusters divided by the total number of identified spectra in non-noise clusters. Spectra are considered incorrectly clustered if their peptide labels deviate from the most frequent peptide label in their clusters, with unidentified spectra not considered.
**Completeness** Completeness measures the fragmentation of spectra corresponding to the same peptide across multiple clusters and is based on the notion of *entropy* in information theory. A clustering result that perfectly satisfies the completeness criterium (value “1”) assigns all PSMs with an identical peptide label to a single cluster. Completeness is computed as one minus the conditional entropy of the cluster distribution given the peptide assignments divided by the maximum reduction in entropy the peptide assignments could provide [34].

Runtime measurements were acquired on a single compute node with two 14-core Intel E5-2680v4 CPUs and 128 GB memory. All tools were allowed to use all available processor cores, except MS-Cluster, which does not have multithreaded capabilities. Memory measurements reflect the peak memory consumption reported by the Moab job scheduler.

### 2.6 Clustering configuration

#### 2.6.1 falcon

Spectrum preprocessing was performed as described in section 2.1. Peaks with intensity below 10 % of the base peak intensity were considered as noise peaks. To convert the spectra to vectors, first virtual vectors with bin width 0.05 *m/z* were created. Next, these vectors were converted to vectors of length 800 using feature hashing.

To match spectra to each other they were first partitioned into 1 *m/z* buckets based on their precursor mass. Next, for each bucket a Voronoi diagram was created consisting of up to 65 536 cells, depending on the number of spectra in the bucket. During querying, at most 32 cells per query were explored. For each spectrum, its 128 nearest neighbors were retrieved via similarity searching. Neighbors whose precursor mass tolerance exceeded 20 ppm were omitted, after which for each spectrum the cosine distances to the maximum 64 nearest neighbors were stored in the sparse pairwise distance matrix.

During DBSCAN clustering dense regions were defined as having minimum two spectra with a cosine distance below 0.05, 0.10, 0.15, 0.20, or 0.25. Clusters were split in a postprocessing step to ensure that pairwise precursor mass differences between cluster members did not exceed 20 ppm.

The clustering result with approximately 1 % incorrectly clustered spectra was obtained with a cosine distance threshold of 0.05.

For the scaling evaluation, subsets of 10, 50, 100, 500, 1000, 1500, 2000, and 2212 randomly selected peak files were used.

#### 2.6.2 MaRaCluster

MaRaCluster (version 1.01.1) [38] was run with a precursor mass tolerance of 20 ppm, and with identical p-value and clustering thresholds −3.0, −5.0, −10.0, −15.0, −20.0, −25.0, or −30.0. Other options were kept at their default values.

The clustering result with approximately 1 % incorrectly clustered spectra was obtained with a p-value and clustering threshold of −30.0.

#### 2.6.3 MS-Cluster

MS-Cluster (version 2.00) [9] was run using its “LTQ_TRYP” model for three rounds of clustering with mixture probability 0.000 01, 0.0001, 0.001, 0.005, 0.01, 0.05, or 0.1. The fragment mass tolerance and precursor mass tolerance were 0.05 Da and 20 ppm, respectively, and precursor charges were read from the input files. Other options were kept at their default values.

The clustering result with approximately 1 % incorrectly clustered spectra was obtained with a mixture probability threshold of 0.000 01.

#### 2.6.4 msCRUSH

msCRUSH (version August 26, 2020) [40] was run using 100 clustering iterations, 15 hash functions per hash table, and cosine similarity threshold 0.3, 0.4, 0.5, 0.6, 0.7, or 0.8. Other options were kept at their default values.

The clustering result with approximately 1 % incorrectly clustered spectra was obtained with a similarity threshold of 0.8.

#### 2.6.5 spectra-cluster

spectra-cluster (version 1.1.2) [12, 13] was run in its “fast mode” for three rounds of clustering with the final clustering threshold 0.999 99, 0.9999, 0.999, 0.99, 0.95, 0.9, or 0.8. The fragment mass tolerance and precursor mass tolerance were 0.05 Da and 20 ppm, respectively, and MGF comment strings were ignored. Other options were kept at their default values.

The clustering result with approximately 1 % incorrectly clustered spectra was obtained with a similarity threshold of 0.999 99.

### 2.7 Code availability

*falcon* was implemented in Python 3.8. Pyteomics (version 4.4.2) [24] was used to read MS/MS spectra in the mzML [26], mzXML, and MGF format. spectrum_utils (version 0.3.5) [3] was used for spectrum preprocessing. Faiss (version 1.7.0) [20] was used for efficient similarity searching. Scikit-Learn (version 0.24.1) [30] was used for DBSCAN clustering, and fastcluster (version 1.1.28) [29] was used for hierarchical clustering. Additional scientific computing was done using NumPy (version 1.20.2) [14], SciPy (version 1.6.2) [36], Numba (version 0.53.1) [23], and Pandas (version 1.2.3) [28]. Data analysis and visualization were performed using Jupyter Notebooks [39], matplotlib (version 3.4.1) [17], and Seaborn (version 0.11.1) [43].

All code is available as open source under the permissive BSD license at https://github.com/bittremieux/falcon. Code used to compare various spectrum clustering tools and to generate the figures presented here is available on GitHub (https://github.com/bittremieux/falcon_notebooks).

## 3 Results

An ideal clustering result groups MS/MS spectra corresponding to distinct peptides in individual, disjoint clusters. The main aspects that influence clustering quality are the spectrum similarity measure and the algorithm that is used to group similar spectra. As various spectrum clustering tools differ in these choices, even when processing identical MS/MS data they will produce cluster assignments with different characteristics. Additionally, the clustering algorithms should exhibit a good computational efficiency to be able to process large amounts of mass spectral data.

The comparison between the different clustering tools in terms of their clustering quality shows that MaRaCluster and spectra-cluster succeed in clustering the highest number of spectra at a comparable rate of incorrectly clustered spectra, while *falcon*, MS-Cluster, and msCRUSH achieve a similar, slightly lower, performance (figure 2A). Notably, the latter tools use the cosine similarity as their spectrum similarity measure. In contrast, MaRaCluster uses a specialized fragment rarity metric to determine spectral similarity [38], while Griss et al. [13] report that replacing the cosine similarity with a probabilistic scoring approach in an updated version of spectra-cluster helped to improve its clustering accuracy.

**Figure 2:**
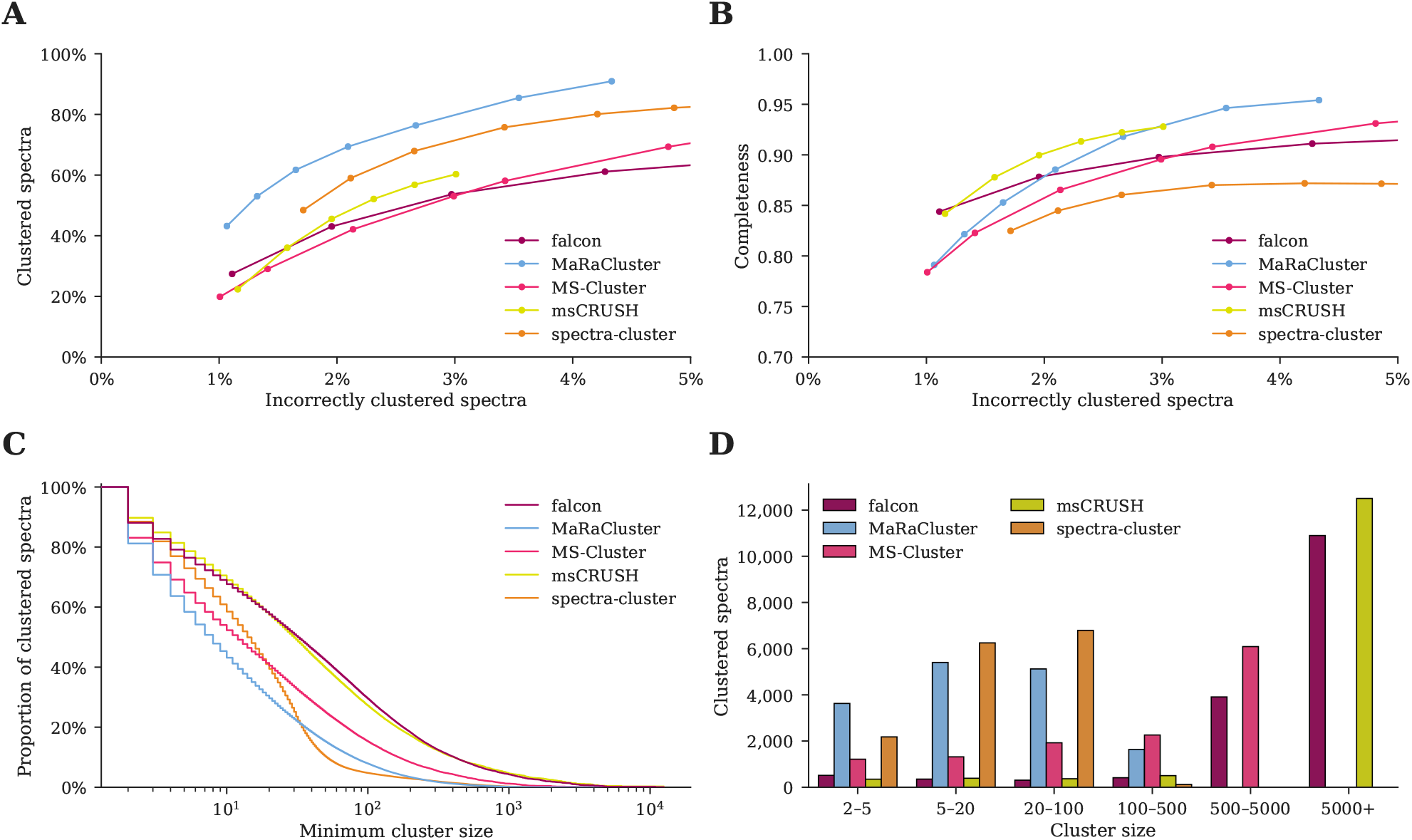
Clustering comparison between *falcon*, MaRaCluster, MS-Cluster, msCRUSH, and spectra-cluster. **(A)** MaRaCluster and spectra-cluster succeed in clustering more spectra than alternative tools at a comparable rate of incorrectly clustered spectra. **(B)** *falcon*, msCRUSH, and MaRaCluster produce a more complete clustering at different rates of incorrectly clustered spectra, grouping spectra corresponding to specific peptides in single clusters. **(C)** Complementary empirical cumulative distribution of the cluster sizes for the clustering results reported in table 1. Although MaRaCluster and spectra-cluster successfully cluster more spectra (less noise points), they predominantly generate clusters consisting of fewer than 100 spectra. In contrast, *falcon* and msCRUSH produce the largest clusters. **(D)** Cluster sizes for the frequently occuring peptide VATVSIPR/2. MaRaCluster and spectra-cluster split spectra corresponding to this peptide into a large number of small clusters consisting of less than 100 spectra each, whereas *falcon* and msCRUSH produce a few large clusters that contain thousands of spectra instead.

Besides generating pure clusters containing spectra corresponding to a single peptide, clusters should also be as complete as possible, i.e. all spectra corresponding to a specific peptide should be grouped in a single cluster. Because each cluster can be represented by a consensus spectrum for subsequent downstream processing, a more complete clustering will result in an increased data reduction by minimizing the generation of redundant cluster representatives. At a low number of incorrectly clustered spectra, *falcon* and msCRUSH achieve the highest completeness, while MaRaCluster outperforms the other clustering tools in terms of completeness at slightly higher numbers of incorrectly clustered spectra (figure 2B). In contrast, spectra-cluster consistently achieves a lower completeness and fails to improve its completeness at increasing numbers of incorrectly clustered spectra.

For clustering results with approximately 1 % incorrectly clustered spectra and minimum cluster size 2 (table 1), MaRaCluster and spectra-cluster predominantly produce small clusters consisting of maximum 100 spectra each (figure 2C). In contrast, *falcon*, msCRUSH, and, to a lesser extent, MS-Cluster produce several larger clusters consisting of 1000 to 10 000 spectra as well. Consequently, even though MaRaCluster and spectra-cluster manage to cluster the highest number of spectra, they produce more small clusters than the alternative tools. Hence, their clustering results will be more fragmented as they do not group all observations of repeatedly occurring peptides together but instead split these spectra across multiple smaller clusters.

**Table 1:**
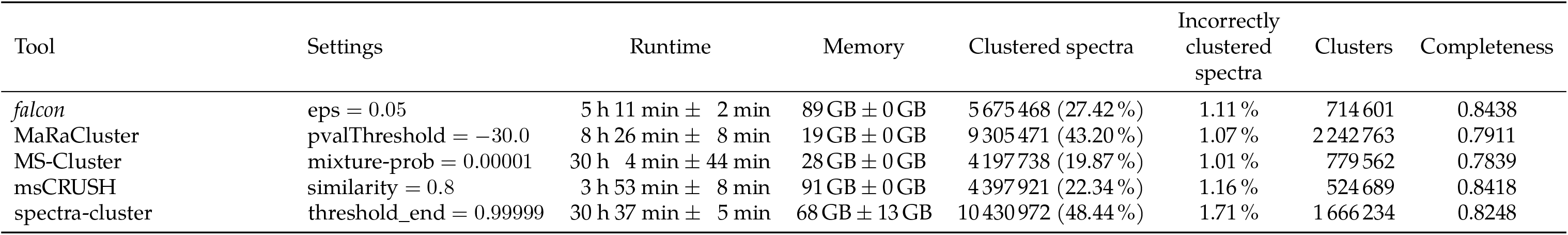
Comparison at an approximate rate of 1 % incorrectly clustered spectra between *falcon*, MaRaCluster, MS-Cluster, msCRUSH, and spectra-cluster in terms of computational requirements and clustering quality. The reported runtimes are the average and standard deviation for three executions of each clustering tool. The reported memory is the average and standard deviation of the peak memory consumption for three executions of each clustering tool. Note that the rate of incorrectly clustered spectra is slightly higher for spectra-cluster, as it was unable to produce highly pure clustering results despite setting very stringent spectrum similarity thresholds.

To better understand the observed differences in completeness of the various clustering approaches, we investigated some clusters manually. For example, VATVSIPR/2, which is part of the pig trypsin contaminant protein P00761, is the most frequently identified peptide in the dataset (observed 18 657 times, figure 2D). Spectra for this peptide are split over 2192 and 1688 separate clusters in the MaRaCluster and spectra-cluster results respectively, and the largest of these clusters only consist of 262 (MaRaCluster) and 121 (spectra-cluster) spectra. In contrast, MS-Cluster groups these spectra into 705 unique clusters, with the largest cluster consisting of 4225 spectra. Finally, *falcon* and msCRUSH split spectra corresponding to this peptide into only 269 and 201 unique clusters respectively, of which the largest cluster produced by both tools alone contains over ten thousand spectra.

As current shotgun proteomics datasets grow ever larger, clustering algorithms need to be scalable to be able to process larger data volumes. For all five clustering tools, settings that are optimized for speed were used to ensure a fair comparison in terms of runtime. For MS-Cluster and spectra-cluster only three rounds of their iterative cluster refinement procedure were used. Additionally, spectra-cluster’s “fast mode” was enabled. For *falcon* a limited number of Voronoi cells per query were examined. For msCRUSH the recommended number of iterations (100) and hash functions (15) for large datasets were used. Runtime measurements include both spectrum clustering and the generation of representative spectra for each cluster. Whereas *falcon* and MS-Cluster can perform the latter functionality directly during clustering, MaRaCluster, msCRUSH, and spectra-cluster require a postprocessing step to export cluster representatives.

msCRUSH exhibited the shortest runtime and was able to process the full dataset consisting of 25 million spectra in only four hours (table 1). Meanwhile, *falcon* processed this data in five hours. Both msCRUSH and *falcon* use advanced algorithmic techniques to reduce the number of pairwise spectrum comparisons that need to be performed. For example, when computing all pairwise spectrum similarities in a brute-force fashion, on average each spectrum in the dataset has to be compared to 1908 other spectra when using a precursor mass tolerance of 20 ppm. In contrast, using its advanced nearest neighbor searching approach, *falcon* on average only had to perform 47 spectrum–spectrum comparisons for each spectrum. Furthermore, the number of neighbors to consider during nearest neighbor searching can be tuned to obtain the desired performance-precision trade-off. Additionally, evaluating different hyperparameters of the density-based clustering step to obtain stricter or looser clusters can be done in only a matter of minutes using a precomputed pairwise distance matrix. In contrast, although MaRaCluster, MS-Cluster, and spectra-cluster use a few simples strategies to avoid having to consider all possible spectrum pairs, such as heuristics based on the number of shared peaks or greedy spectrum merging strategies, depending on their settings, these tools can be considerably slower.

To further test how the *falcon* runtime scales in terms of its input data, subsets of the human draft proteome dataset consisting of 10 to 2212 randomly selected peak files were clustered (figure 3). Whereas theoretically the number of pairwise spectrum comparisons scales quadratically in terms of the number of processed spectra, the *falcon* runtime exhibits only a linear scaling. This computational efficiency is an essential requirement to be able to expediently process the ever increasing data volumes generated during mass spectrometry proteomics experiments, and it indicates that clustering of large data volumes in public data repositories can be feasible.

**Figure 3:**
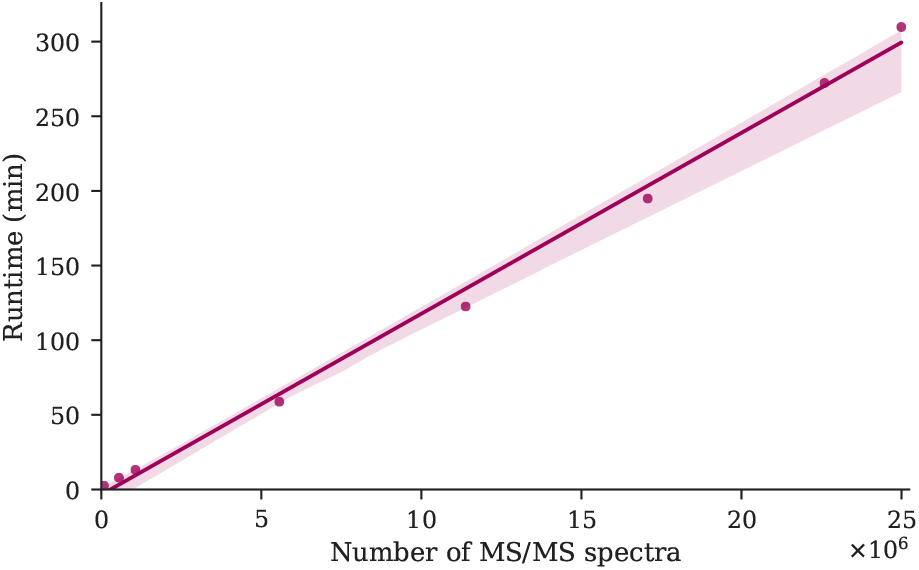
The *falcon* runtime for an increasing number of spectra to be clustered shows linear scalability in terms of the input size.

## 4 Conclusion

Here we have introduced the *falcon* spectrum clustering tool. *falcon* uses various advanced algorithmic approaches, such as feature hashing to vectorize high-dimensional spectra and fast nearest neighbor searching. It exhibits a high computational efficiency and outperforms most alternative spectrum clustering tools in terms of runtime while generating clusters of a comparable quality. As such, *falcon* is ideally suited to process the ever increasing amounts of mass spectral data generated during mass spectrometry experiments.

Although *falcon* succeeds in clustering a similar number of spectra as MS-Cluster and msCRUSH at a comparable rate of incorrectly clustered spectra, these three clustering tools are outperformed by MaRaCluster and spectra-cluster in terms of the number of clustered spectra. Notably, *falcon*, MS-Cluster, and msCRUSH compare spectra to each other using the cosine similarity, whereas MaRaCluster and spectra-cluster use more advanced scoring approaches. The cosine similarity is a commonly used similarity measure [37]. Some of its advantages are that it can accurately capture spectrum similarity, it is easy to implement and interpret, and it is fast to evaluate. Consequently, the cosine similarity is a highly competitive and ubiquitous baseline method. However, alternative spectrum similarity methods, such as spectra-cluster’s probabilistic score [13], MaRaCluster’s fragment rarity score [38], or similarity measures derived from machine learning [15, 27] might be able to more sensitively capture spectrum similarity. Additionally, it is important to evaluate how MS/MS spectrum preprocessing [32] influences various similarity measures.

We have evaluated several state-of-the-art spectrum clustering tools based on characteristics of the clusters they produce, such as the number of incorrectly clustered spectra, cluster completeness, and cluster size, rather than indirectly evaluating their performance on a downstream task. Myriad applications of spectrum clustering exists, such as the compilation of spectral libraries [12, 42], molecular networking [41, 44], and as a data reduction technique prior to computationally intensive analyses, such as open modification searching [5]. The performance of these applications does not only depend on the spectrum clustering quality, but also the strategy used to form cluster representatives (e.g., selecting the medoid spectrum with minimum average distance to all cluster members, compiling a consensus spectrum by merging all cluster members in a specific fashion, or alternative methods) and the settings of other tools in the bioinformatics workflow. As demonstrated in our evaluation of five spectrum clustering tools, alternative tools exhibit different performance characteristics. As such, the optimal tool to use will likely depend on the practitioners’ downstream task.

Among the five clustering tools evaluated, at a low number of incorrectly clustered spectra, *falcon* generates a highly complete clustering result. As such, it will achieve a large reduction in data volume when representing the clustered data by their consensus spectra, which can be especially relevant when spectrum clustering is used as a preprocessing step prior to a subsequent computationally intensive analysis. Additionally, the reduction of redundant information can facilitate downstream interpretation, for example, by avoiding uninformative nodes and edges during molecular networking.

A complicating factor for the evaluation of spectrum clustering tools is how chimeric spectra are handled. As these spectra contain fragments of multiple distinct ions, they cannot be unambiguously assigned to only a single cluster. Additionally, chimeric spectra can potentially bridge clusters corresponding to different peptides, incorrectly producing a single, heterogeneous, cluster. Although *falcon* does not explicitly guard against this event, its competitive performance compared to alternative spectrum clustering tools in terms of correctly clustered spectra indicates that it does not unduly suffer from the presence of chimeric spectra and is able to handle them in a robust fashion. Nevertheless, the full extent of the effect of chimeric spectra on spectrum clustering and identification currently still remains an important open research question.

*falcon* is freely available as open source under the permissive BSD license. The source code and instructions can be found at https://github.com/bittremieux/falcon.

## Acknowledgments

We want to thank Kilian Maes and Laurent Gatto for fruitful discussions about the *falcon* performance.

W.B. is a postdoctoral researcher of the Research Foundation – Flanders (12W0418N). The resources and services used in this work were provided by the VSC (Flemish Supercomputer Center), funded by the Research Foundation – Flanders (FWO) and the Flemish Government. This work was supported in part by National Institutes of Health award U19 AG063744-01.

